# Quantifying the relationship between spreading depolarization and perivascular cerebrospinal fluid flow

**DOI:** 10.1101/2023.04.10.536161

**Authors:** Saikat Mukherjee, Mahsa Mirzaee, Jeffrey Tithof

## Abstract

Recent studies have linked spreading depolarization (SD, an electro-chemical wave in the brain following stroke, migraine, traumatic brain injury, and more) with increase in cerebrospinal fluid (CSF) flow through the perivascular spaces (PVSs, annular channels lining the brain vasculature). We develop a novel computational model that couples SD and CSF flow. We first use high order numerical simulations to solve a system of physiologically realistic reaction-diffusion equations which govern the spatiotemporal dynamics of ions in the extracellular and intracellular spaces of the brain cortex during SD. We then couple the SD wave with a 1D CSF flow model that captures the change in cross-sectional area, pressure, and volume flow rate through the PVSs. The coupling is modelled using an empirical relationship between the excess potassium ion concentration in the extracellular space following SD and the vessel radius. We find that the CSF volumetric flow rate depends intricately on the length and width of the PVS, as well as the vessel radius and the angle of incidence of the SD wave. We derive analytical expressions for pressure and volumetric flow rates of CSF through the PVS for a given SD wave and quantify CSF flow variations when two SD waves collide. Our numerical approach is very general and could be extended in the future to obtain novel, quantitative insights into how CSF flow in the brain couples with slow waves, functional hyperemia, seizures, or externally applied neural stimulations.

## 1 Introduction

The human brain contains about 80% water in different compartments including blood, cerebrospinal fluid (CSF) in the ventricles, and interstitial fluid (ISF) in the extracellular space (ECS)^1^. The recent glymphatic hypothesis suggests that the water in the brain is dynamic, with exchange of CSF and ISF to help clear metabolic waste. Notably, CSF has been shown to flow through annular spaces that line the brain vasculature, known as perivascular spaces (PVSs)^2–7^. Although, the exact nature of exchange between CSF in the PVS and ISF in the brain tissue is still debated^8^, it is noteworthy that even a laminar and viscous fluid flow may significantly enhance transport since the diffusion coefficient of waste protein molecules is relatively small (leading to a Péclet number Pe ≥ 1 for large molecules like amyloid-*β* ^9, 10^).

The ISF in the ECS is primarily a diluted salt solution that acts as an ionic reservoir consisting of Na^+^, K^+^, Ca^2+^, and Cl^−^. The ISF promotes important neuronal functions like regulating the pH level, maintaining the resting potential of neurons, and enabling action potential propagation^11^. The neurons, glia, and vasculature in the brain tissue are bathed by the ISF which pervades the ECS and constitutes about 20% of the total brain volume^12^. The presence of substantial amounts of freely floating charged ions and water in the brain thus lays the perfect grounds for rich chemo-hydrodynamic interactions between ionic fluxes and the dynamics of CSF and ISF.

Maintaining ionic fluxes in the ECS at homeostasis requires a substantial amount of metabolic energy. Indeed, the human brain accounts for about 20 − 25% of the total metabolism even though it constitutes only about 2% of the body mass^13^. The ion channels and pumps in the neuronal cells use this energy to transport ions between the intracellular space (ICS) and ECS. Ionic fluxes can interact with CSF and ISF in complicated ways exemplified by neurovascular coupling, generation of osmotic pressure gradients, and chemical reaction-diffusion waves. Under physiological conditions, neurovascular coupling causes local changes in the cerebral blood flow which can lead to changes in the CSF flow, as demonstrated in functional hyperemia and in slow wave oscillations during sleep^14–16^.

In some acute neurological conditions, the ionic equilibrium may become compromised resulting in spreading depolarization (SD). SD is a reaction-diffusion wave resulting from redistribution of ions in the ECS and ICS during stroke, traumatic brain injury, migraine, cardiac arrest, and haemorrhage^8,17–20^. First observed by Leão^21^, SD is characterized by an influx of ions like Na^+^ and Ca^2+^ into the cells and an outflow of K^+^. Under physiological conditions, a single neuron maintains a potential of about 70 mV due to the ion gradients maintained across the cell membrane. Depolarization occurs when the ionic redistribution increases the resting potential to a value of approximately − 10 mV^17,22^. SD-induced ionic gradients across the cell membrane also lead to swelling of the neurons because of water influx into the cell from the ECS due to osmotic stress. Additionally, the excess K^+^ in the ECS leads to changes in the lumen radius of the arteries. The excess ions in the cortex then spread due to diffusion along the cortical surface leaving behind a trail of swelled up depolarized neurons and constricted vasculature in their wake. The resulting SD wave has a typical speed of 𝒪 (1) mm/min^23–25^.

Previous studies of chemical reaction-diffusion waves in an imposed fluid flow have shown that reaction-diffusion waves can disrupt complex fluid flow through different forms of coupling^26,27^. There are several mechanisms by which SD may disrupt the glymphatic system. A recent study found that SD due to induced ischemic stroke in mice resulted in large CSF influx into the brain, a novel discovery indicating CSF is the earliest contributor to brain edema (water accumulation in tissue) following stroke^8^. Using two-photon imaging and numerical modeling, it was found that SD-induced vasoconstriction of penetrating arterioles induced a large bulk flow of CSF by increasing the effective size of the surrounding PVSs. SD is also linked to edema and blood-brain-barrier compromise following cardiac arrest due to transmembrane ionic gradients, which cause substantial water influx and tissue destruction^28^. In addition to ischemia, SD also occurs during chronic migraine^18,19^ and traumatic brain injury^29,30^. Moreover, apart from vasoconstriction and osmotic stresses, the shrinkage of ECS volume during SD may also disrupt CSF-ISF exchange and transport in the interstitial spaces^31–34^.

Despite great progress in experimental techniques, quantifying the precise mechanisms coupling SD and CSF and/or ISF dynamics *in vivo* is difficult because of spatiotemporal limitations in imaging. Physiologically realistic numerical simulations offer an approach that sidesteps this limitation. In this work we numerically simulate one of the several important mechanisms that couple SD and the glymphatic system as outlined above. We focus on the chemo-mechanical coupling during SD that leads to vasoconstriction/vasodilation in pial and penetrating arteries resulting in CSF influx to the brain. We follow an empirical relationship between the K^+^ concentration and the arterial lumen radius^35–37^. It is important to note that the excess K^+^ during SD in the ECS is first transported via diffusion to the PVS. The increased K^+^ concentration in the PVS then causes a change in the polarization of the smooth muscle cells lining the arterial lumen, leading to vasoconstriction or vasodilation^36^. This excess K^+^ in the PVS is depleted after the SD wave has passed and is also perhaps removed by spatial potassium buffering^38^. It is useful to note that the neurovascular coupling during functional hyperemia and slow wave oscillations during sleep also share similarity with the mechanism outlined above, albeit the K^+^ concentrations are lower. On the other end of the spectrum, seizures can cause severe and sustained vasoconstrictions^39^ which share mechanistic similarities with SD^40^. Additionally, recent studies have shown that externally applied nerve stimulation can also lead to increasing CSF tracer penetration into the brain, possibly due to similar hemodynamic alterations^41^. Hence, novel insights into CSF transport due to changes in arterial diameter have far-reaching implications beyond SD alone.

To model the SD wave we implement a 13-component reaction-diffusion system of equations, motivated by physiological ionic dynamics, which yields a traveling SD wave in one spatial dimension. To model the CSF flow in the PVS we solve a system of one-dimensional (1D) Navier-Stokes equations which quantify the changes in volumetric flow rate and pressure as a function of the change in PVS area during SD. A 3D or 2D representation of CSF flow in the PVS becomes computationally prohibitive because of the need to resolve the disparate length scales of the problem. The wavelength of a typical spreading depolarization is *λ* = 𝒪(1) mm while the PVS width is *δ*_*r*_ = 𝒪 (10^−3^) mm, which makes the problem computationally difficult because of the large aspect ratio (*λ*/*δ*_*r*_ = 𝒪(10^3^)). Nevertheless, full simulations in idealized geometries and image-based geometries have elucidated the pressure and fluid flow distribution in the PVS^42,43^. Simulations in 1D offer the advantage of computational efficiency with a relatively small compromise on the accuracy of average volumetric flow rate and pressure magnitudes^44^. Moreover, 1D simulation of CSF is also motivated by the substantial volume of work on modeling blood flow in vascular networks^45^. The numerical approach we implement is very flexible and can be easily adapted to quantify CSF flow in the brain resulting from slow waves, functional hyperemia, seizures, different forms of SD, or externally applied nerve stimulations.

This article is structured as follows. We first describe our approach by stating the governing equations of SD and CSF flow, as well as our numerical implementation. We then detail our results, where we explore the coupling of the SD wave and CSF flow by systematically varying multiple parameters, which include the arterial radius, the PVS width, the PVS length, and the angle of incidence of the SD wave on the PVS. Additionally, we also explore the variation of PVS area, volumetric flow rate of CSF, and pressure changes following a collision of SD waves. We end with concluding remarks.

### 2 Approach

We first describe the domain in which we perform the numerical simulations. Our numerical domain consists of a single branch of an artery of variable length *L*_PVS_ sheathed by an annular PVS. The width of the PVS is *δ*_*r*_ = *r*_2_ − *r*_1_, where *r*_1_ is the lumen radius and *r*_2_ is the radius of the outer wall of the PVS. We then model a traveling SD wave incident on the PVS at a variable angle 0 ≤ *θ* ≤ *π*/2. Figure 1 shows our numerical domain. Our modelling approach is to couple the Yao-Huang-Miura model (YHM) of spreading depolarization^46^ with a model of 1D fluid flow through a perivascular space.

**Figure 1.**
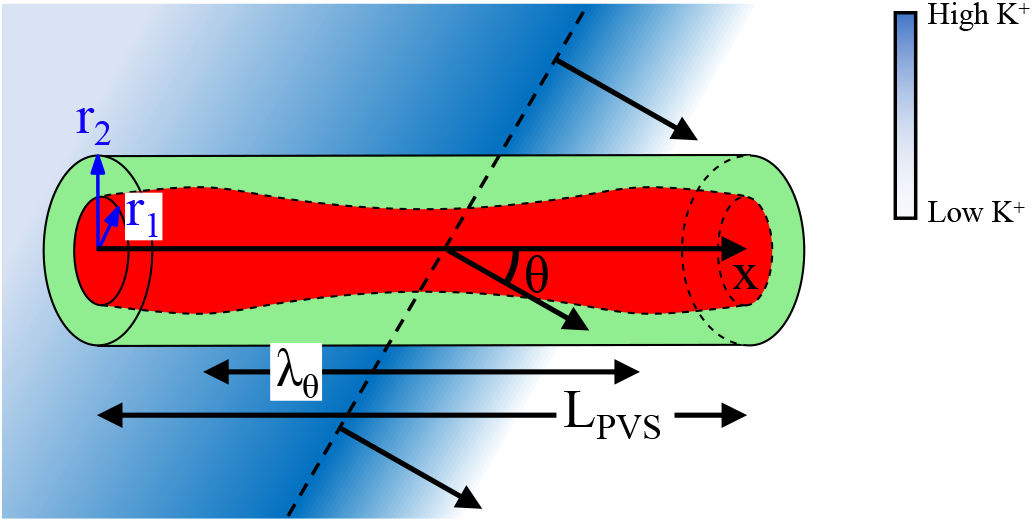
The schematic of our computational domain. A single arterial segment is shown in red, which is ensheathed by PVS shown in green. The extracellular potassium concentration, associated with the SD wave, is shown in blue. The dashed line indicates the maximum K^+^ concentration, and the three perpendicular arrows indicate the direction of the wave propagation. The figure is not drawn to scale.

### 2.1 Spreading depolarization

Mathematical modeling approaches to SD usually fall under two categories, qualitative and physiological. Qualitative approaches model the excitatory-inhibitory dynamics associated with SD using phenomenological reaction-diffusion models. The traveling wave in these models constitute a bifurcation from the base state of the system due to a change in a control parameter^18,47,48^. While these models recreate the 2D spatial dynamics exhibited by SD, like spiral and stationary waves, it can be difficult to connect the qualitative variables with ionic dynamics. On the other hand, the physiological approach involves solving coupled systems of reaction-diffusion equation of physiological ionic transport in the brain including diffusion in the ECS, intracellular dynamics, electro-diffusion, as well as ECS shrinkage due to osmotic water flow into the cells^23,34,46,49,50^. The limitations of the physiological models are that the governing equations are usually multi-variable and computationally cumbersome. Additionally, physiological models are typically implemented in 1D because of computational expense, and they may be unable to recreate the 2D spatial patterns exhibited by SD waves; recently however, a physiological model based on electro-diffusion has been proposed that yields spiral waves when simulated in 2D^51^. Below we discuss our modeling approach of SD where we have implemented the physiologically-realistic YHM model.

YHM is a comprehensive model of spreading depolarization that takes into account the ionic transport across the neuronal membrane, cross-membrane currents, and membrane potential variation in the intracellular neuronal space. These mechanisms are coupled with diffusion of ions in the extracellular space^46^. The model uses the technique outlined in Kager *et al*.^52^ to quantify the intracellular dynamics of ions and currents across the neuronal membranes. The intracellular ionic dynamics are coupled with the continuum modelling approach developed by Tuckwell & Miura^49^ in the extracellular space. It is assumed that the ICS and the ECS share the same overlapping continuum space. We use the basic YHM model which captures the dynamics of Na^+^ and K^+^ ions; however, more complicated variations exist which incorporate cell swelling and Cl^−^ ions.

The basic formulation of the YHM model is as follows:

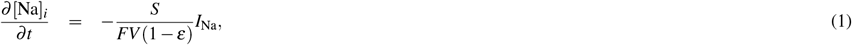

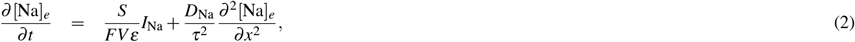

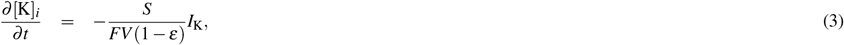

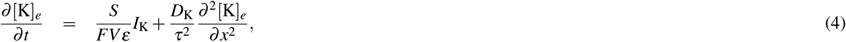

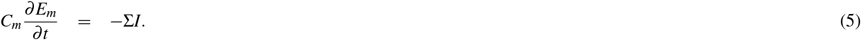

Here time is expressed in the units of ms and space is expressed in the units of cm. In Eqs. (1-5), *i* represents the intracellular space, *e* represents extracellular space, *S* =1.6 × 10^−5^ cm^2^ is the neuronal surface area, *F* =96. × 485 coulomb*/*mM is the Faraday constant, and *ε* =*V*_*e*_/*V* is the extracellular volume fraction. For the present study we have used *ε* = 0.13 which implies that cells occupy 87% of the brain interstitium (*V* is the total volume such that *V* = *V*_*i*_/(1−*ε*), where *V*_*i*_ = 2.2 × 10^−9^ mm^3^ is the typical value of neuronal volume). *E*_*m*_ is the neuronal membrane potential. Other relevant parameters are the neuronal membrane capacitance *C*_*m*_ = 7.5 × 10^−7^ s/Ω cm^2^, the diffusion coefficient of sodium and potassium ions *D*_Na_ = 1.33 × 10^−5^ cm^2^/s and *D*_K_ = 1.96 × 10^−5^ cm^2^/s^50,53^, and the tortuosity *τ* of the extracellular space. Since the brain interstitium can be considered as a porous medium, the diffusion of ions in the extracellular space is hindered when compared to an open medium; *τ* quantifies the hindrance of diffusive spreading in a porous medium. In the volume-averaged formulation of the transport equations in a porous medium, the diffusion coefficient is scaled by *τ*^2^ and the porosity *ε* is incorporated in the source terms as implemented in Eq. (2) and Eq. (4)^12^. We have used *τ* = 1.55 which agrees with experimental measurements of ion diffusion in a rat cerebellum^54^. Tortuosity depends on the pore structure of the interstitium and can vary between species; however, the relationship between SD propagation speed *v* and *τ* is expected to be *v* ∝ 1*/τ*, since reaction-diffusion wave speed scales with the diffusion coefficient as *v* ∝ 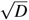 ^55, 56^.

The total cross-membrane ionic currents due to the movement of Na^+^ and K^+^ ions across the membrane is given by the sum of sodium and potassium currents, Σ*I* = *I*_Na_ + *I*_K_ + *I*_leak_^46^, where *I*_leak_ is the general leak current given by *I*_leak_ = *g*_*HH*_(*E*_*m*_ + 70). The sodium and potassium currents consist of active currents which are modelled by Goldman-Hodgkin-Katz (GHK) equations, and passive (or leak) currents which are modelled by Hodgkin-Huxley (HH) equations. The sodium current *I*_Na_ = *I*_Na,*P*_ + *I*_Na,*T*_ + *I*_Na,leak_ + *I*_Na,pump_, consists of active currents, namely fast sodium current (*I*_Na,*T*_) and persistent sodium current (*I*_Na,*P*_). The potassium current *I*_K_ = *I*_K,DR_ + *I*_K,A_ + *I*_K,leak_ + *I*_K,pump_ consists of active current, namely the potassium delayed rectifier current (*I*_K,DR_) and the transient potassium current (*I*_K,A_). The GHK and HH equations modelling the active and leak currents respectively are,

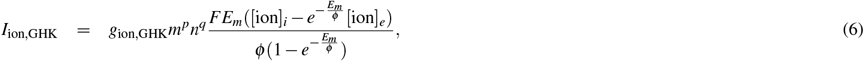

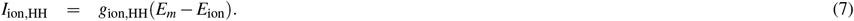

Here 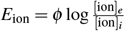 is the Nernst potential while *ϕ* = *RT*/*F* is a parameter where *R* = 8.31 mV·C/(mM·K) is the universal gas constant and *T* = 298 K is the absolute temperature. Additionally, *g*_ion,HH_ is the conductance amplitude for passive currents whereas *g*_ion,GHK_ is the product of the neuronal membrane potential and conductance amplitude for the active currents. The active currents are voltage gated by specific gating proteins which allow for the transport of ions across the neuronal membrane depending on the membrane potential. The gating variables can be expressed as,

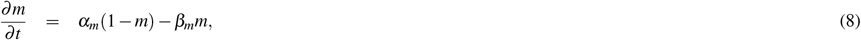

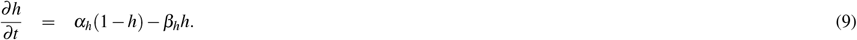

Here *m* and *h* are first order activation and inactivation gates respectively. Rate constants *α*_*m*_(*E*_*m*_), *β*_*m*_(*E*_*m*_), *α*_*h*_(*E*_*m*_), and *β*_*h*_(*E*_*m*_) are functions of the membrane potential. Finally, the total ionic currents also consist of the pump currents, *I*_K,pump_ = 3*I*_pump_ and 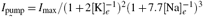 is the general potassium-sodium exchange pump current and *I*_max_ = 0.013 *mA/*cm^2^. Supplementary Table S(1), S(2), and S(3) in the supplementary information lists the values of conductances, currents, and rate constants used in the simulation. It is useful to note that electroneutrality is not enforced in the present model. Although electroneutrality is maintained in the brain under normal conditions, it is unclear whether it is maintained during SD^23^. Our modeling approach follows previous models of SD where only cations are considered^46,49,52^. However, electroneutrality should also be enforced if the dynamics of anions like Cl^−^ in addition to cations are studied in the future using this model^46^. In addition, electro-diffusion models can also be explored in the future^51^.

To solve Eqs. (1-5) and Eqs. (8-9) we use a fourth order accurate Runge-Kutta scheme in time, and second order accurate finite differences in space. The numerical solver is written in Fortran. We use a time step of Δ*t* = 5 × 10^−3^ ms, and a grid spacing of Δ*x* = 0.01 mm. Since there are no exact solutions, we have used the wave speed *v* at the finest grid resolution (Δ*x* = 0.002 mm) possible for this time step to compare our results. A plot of error as a function of the grid resolution is included in Supplementary Fig. S(2) in the supplementary information.

The initial condition we use for the extracellular potassium concentration is in the form of a Gaussian given by,

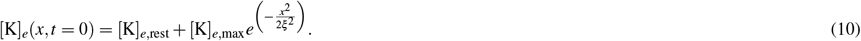

This initial condition models a spike in potassium concentration at a localized region in space of width *ξ* = 0.5 × 10^−2^ mm, which is the case when KCl is applied to instigate SD waves in experiments^57^. The initial spike is of magnitude [K]_*e*,max_ = 50 mM. The initial resting values of all the ions and gating variables are noted in Supplementary Table S(3) of the supplementary information. The boundary condition for [K]_*e*_ and [Na]_*e*_ are no-flux at *x* = 0 and *x* = *L*_SD_, where *L*_SD_ = 6 mm is the spatial extent that we simulate the SD wave.

### 2.2 1D Navier-Stokes equation

Solving Eq. (1-9) results in a traveling wave of [K]_*e*_, [Na]_*e*_ and *E*_*m*_, in the positive *x*-direction. We next couple the spreading wave with the 1D Navier-Stokes equation. The coupling occurs due to a change in radius of the artery, induced by a spike in the [K]_*e*_ concentration due to the SD wave. The traveling wave of [K]_*e*_ leads to a spatial change in radius of the arteries which travels with the same velocity as the SD wave, when the arteries are aligned with the direction of wave propagation. We note that the radius of the artery forms the inner boundary of the PVS, *r*_1_. We thus expect a change in the fluid dynamics of the CSF in the PVS, which can be induced by the SD wave. The conservation of mass and momentum for a 1D section of PVS subjected to a SD wave can be written as,

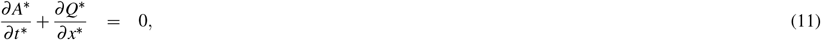

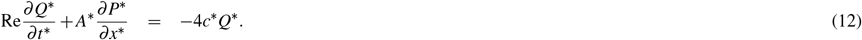

Here *A*^∗^, *Q*^∗^, and *P*^∗^ are the nondimensional area of the perivascular space (PVS), average volumetric flow rate, and average pressure respectively. We use the symbol to denote nondimensional variables throughout the text.

Equations (11-12) are derived using a similar approach to Ref.^58^. In this approach, the governing equations of fluid flow are first cast using cylindrical coordinates in an annulus, assuming axisymmetric flow. The underlying assumption we then use is the lubrication approximation which is valid when the ratio of the length scales in the radial direction to the axial direction is small. In our setup, *δ*_*r*_/*λ* << 1. This approximation allows us to neglect the variation of pressure in the radial direction 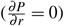. The equations are then averaged in the radial direction and the variation of the arterial radius with time is incorporated. We use the steady state Poiseuille flow profile in an annulus to further reduce the radially averaged governing equations. The equations are nondimensionalized using a length scale of PVS width *δ*_*r*_ = *r*_2_ − *r*_1_ in the radial direction and a length scale *λ* (the wavelength of the SD wave) in the axial direction. The time scale used for nondimensionalization is *T* = *λ*/*v*, where *v* is the wave speed. *T* is defined as the wave period. We note that the frequency of the wave is *f* = 1/*T* = *v*/*λ* . We have included a detailed derivation of Eqs. (11)-(12) in the supplementary information. The variable *c*^∗^ is obtained from the Poiseuille flow profile through a concentric circular annulus, and its dimensional value is

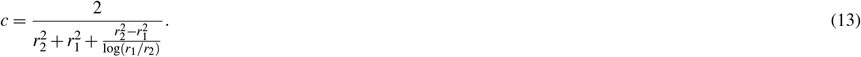

Although we assume the PVS to be a concentric circular annulus, it is important to point out that PVSs surrounding the pial arteries in mice have been shown to have varying eccentricity and ellipticity which can influence the hydraulic resistance through the PVSs^67,68^. Indeed, the hydraulic resistance has been found to be lower than a concentric circular annulus and the flow has been shown to be three-dimensional^67,68^. Additionally, we have assumed all the PVSs surrounding the pial and the penetrating arteries as open medium. Recent studies have indicated that the pial PVSs are open; however, it is unknown as to whether the penetrating PVSs are porous^5,69^. It would be interesting in the future to explore the coupling of SD and CSF flow through porous PVSs.

It is useful to point out that using the above mentioned non-dimensionalization, we retrieve an area scale of a 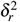 volumetric flow rate scale 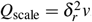, and a pressure scale 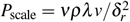, where ν is the coefficient of kinematic viscosity and *ρ* is the density. The Reynolds number Re in Eq. (12) is the ratio of inertial to viscous forces. Using the averaging approach outlined above, the Reynolds number can be written as

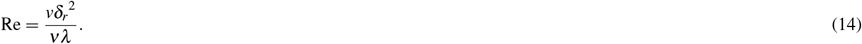

Typically, Re is very small (*O*(10^−3^)) for this problem. Eq. (11-12) are solved using a finite difference approach that implements a predictor-corrector algorithm which is 4th order accurate in time and 2nd order accurate in space. The initial condition is *Q*^∗^ = 0 and *P*^∗^ = 0 inside the domain. At time *t*^∗^ = 0, a deformation of the cross-sectional area *A*^∗^ is prescribed based on the SD-induced deformation of *r*_1_. We assume the outer wall of the PVS *r*_2_ is stationary. The boundary conditions for pressure is *P*^∗^ = 0 at *x* = 0 and *x* = *L*_PVS_ respectively, where *L*_PVS_ is the length of the simulated PVS domain. We have conducted extensive spatial and temporal resolution tests to verify our numerical approach (for more details the reader is directed to the supplementary information).

### 2.3 Relationship between arterial lumen radius and extracellular potassium ion concentration

The lumen radius of the arteries (*r*_1_) changes as a function of the excess potassium concentration during SD. To quantify this change in our model, we have used an empirical relation that relates the change in *r*_1_ to [K]_*e*_ from experimentally realistic numerical models of neurovascular coupling^35,36^. The equation, adapted for our specific problem, relates the normalized lumen radius 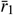 and [K]_*e*_ as

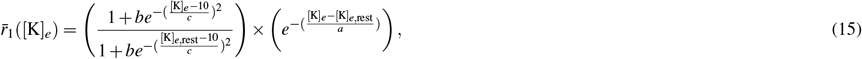

where *a* = 50 mM, *b* = 0.18, *c* = 3 mM, and the resting value of potassium ion concentration [K]_*e*,rest_ = 3.86 mM. Equation (15) is the primary relationship which couples the SD wave to the fluid flow through the PVSs in our model. The first term inside the parenthesis before the multiplication sign on the right hand side of Eq. (15) is the dilation response of 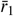 for small values of potassium ion concentrations. The second term inside the parenthesis after the multiplication sign is the constriction response for larger values of [K]_*e*_. As [K]_*e*_ increases, 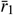 dilates from the base value of 1 to 1.16 for [K]_*e*_ *<* 14 mM before constricting with further increase in potassium ion concentration. A plot of 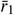 as a function of [K]_*e*_ is included in Supplementary Fig. S(1) in the supplementary information.

## 3 . Results

We first plot the spatiotemporal variations of the ionic concentrations and membrane potential as the SD wave propagates in the cortex in Fig. 2. Figure 2 (a) shows the variation of potassium ion concentration ([K]_*e*_) as a function of the spatial extent of the extracellular space at different instances of time. The initial spike in potassium concentration travels to the right with a velocity of *v* = 4.97 mm/min which we find by tracking the *x*-location of [K]_*e*_ = 20 mM. We quantify the wavelength of the SD wave by determining the maximum width of the wave for [K]_*e*_ > 4 mM, which yields *λ* = 2.5 mm. It is useful to note that the initial resting value of potassium is [K]_*e*_ = 3.86 mM. Figure 2 (b) shows the variation of the normalized inner radius of the PVS, 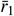, as a function of *x* at the same instances of time shown in Fig. 2 (a). Consistent with Eq. (15), we find in Fig. 2 (a-b), that for [K]_*e*_ < 14 mM, there is a small magnitude of vasodilation (almost 20%). However, the arteries undergo vasoconstriction for [K]_*e*_ > 14 mM by about 80%.

**Figure 2.**
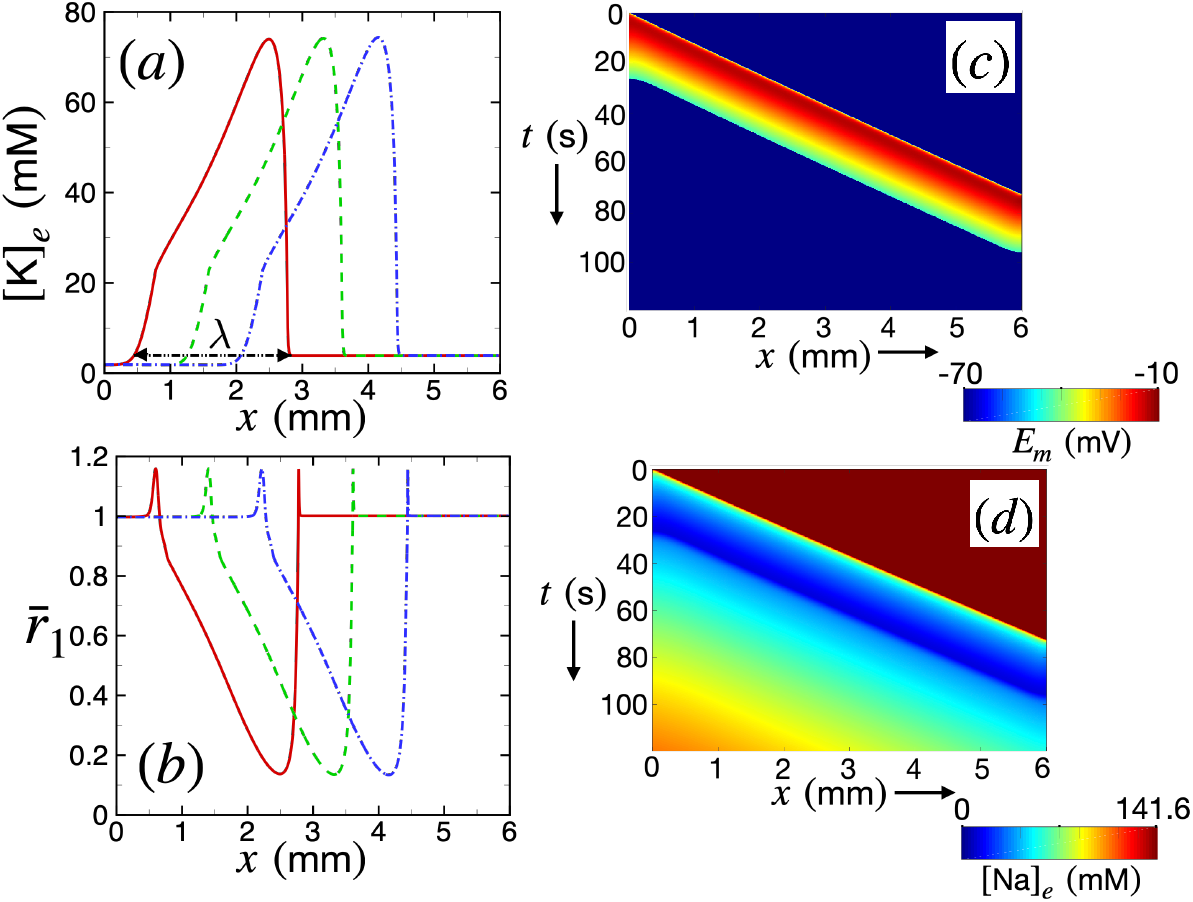
The variation of ionic concentrations, arterial radius, and membrane potential following spreading depolarization. (a) The variation of extracellular potassium ion [K]_*e*_ (mM) as a function of *x* at three different instances of time, *t* = 34 s (red, solid curve), 44 s (green, dashed curve), and 54 s (blue, dashed-dotted curve), respectively. (b) The variation of normalized inner radius 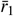 as a function of *x* at the same instances of time as (a). (c) Spatiotemporal variation of membrane potential *E*_*m*_ (mV) shown using a space-time plot. The color contours of *E*_*m*_ are shown. (d) Space-time plot showing the spatiotemporal variation of the concentration of extracellular sodium ions [Na]_*e*_.

Figure 2 (c) shows a space-time plot of membrane potential *E*_*m*_. The typical value of the ionic potential of neurons is −70 mV. The neuronal membrane potential increases to − 10 mV due to SD. As the wave propagates forward, new cells are depolarized, which then revert back to the base state after the wave has passed through. This is evident from the space-time plot where the leading portion of the wave drives a substantial depolarization (red). The trailing portion of the wave is where the neurons gradually start repolarizing to base state (green and yellow). The inverse of the slope of the contours in the space-time plot is equal to the wave velocity *v*.

Figure 2 (d) shows the space-time plot of extracellular sodium concentration [Na]_*e*_. The base state value of sodium concentration in the extracellular spaces is 141.6 mM. Spreading depolarization causes a large influx of sodium ions into the neurons leading to a decrease in extracellular concentration. This is evident from the figure where the leading portion of the wave is immediately followed by a substantial decrease of [Na]_*e*_. As the wave passes through, the trailing cells gradually regain their base state as the [Na]_*e*_ is restored.

We next show the coupling of the SD wave with CSF flow in the PVS in Fig. 3, where we have considered a PVS segment of length equal to the 2.5 times the wavelength of the SD wave, *L*_PVS_ = 2.5*λ* = 6 mm. We have also considered a baseline arterial radius of *r*_1_ = 23 *μ*m, and a PVS width of *δ*_*r*_ = 12.63*μ*m, which match with typical values for pial arteries in the murine brain^9^. Additionally, we have considered a PVS segment which is aligned with the direction of the wave propagation (i.e., *θ* = 0).

**Figure 3.**
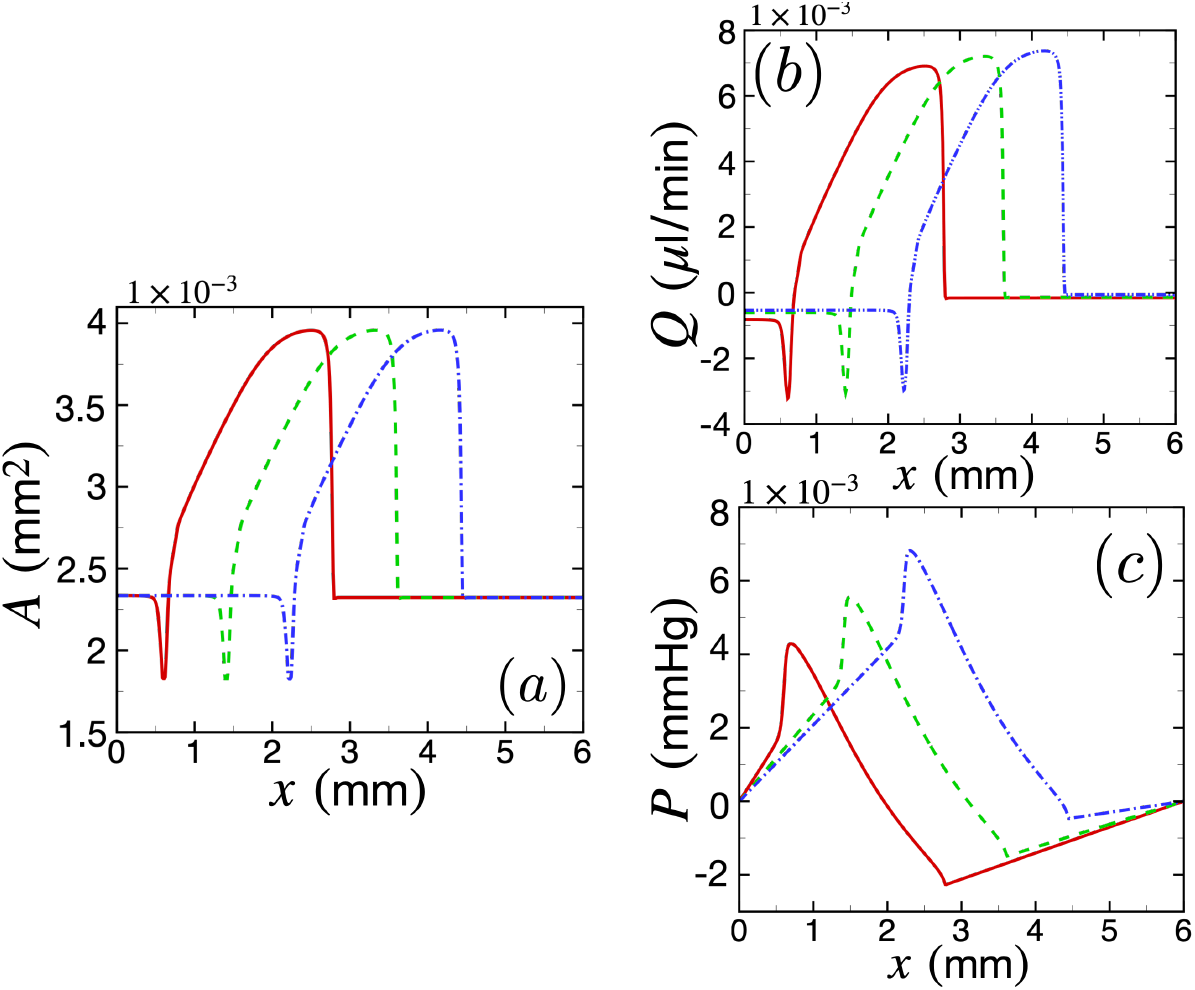
The variation of the PVS area, CSF volume flow rate, and pressure following spreading depolarization. (a) The variation of the area of the PVS *A* as a function of *x* at three instances of time *t* = 34 s (red, solid curve), 44 s (green, dashed curve), and 54 s (blue, dashed-dotted curve). (b) The variation of CSF volume flow rate *Q* as a function of *x* using the same plotting convention as (a). (c) The variation of pressure *P* as a function of *x* using the same convention as (a-b). The relevant parameters are *r*_1_ = 23 *μ*m, *L*_PVS_ = 6 mm, *θ* = 0, and Γ = 1.4.

In Fig. 3 (a), we have plotted the dimensional PVS area *A* as a function of *x* at three different instances of time as the SD wave propagates, using the same convention as Fig. 2 (a). For coupling the SD with CSF flow, we have filtered the radius data to smooth out the sharp peak in the leading edge of the wave (around *x* ≈ 3 mm for the red curve in Fig. 2 (b)). We found that resolving the sharp peak at the leading edge requires extremely fine spatial resolutions that make our simulations computationally expensive, which is why we have smoothed out that peak. We have conducted sufficient tests to establish that smoothing out the peak does not affect our results substantially (a detailed discussion can be found in Sec. 5 of the supplementary information). Since at lower concentrations of [K]_*e*_, arteries undergo vasodilation, the PVS area undergoes a small constriction from the baseline value of 2.34 × 10^−3^ mm^2^ to 1.83 × 10^−3^ mm^2^ or by 27% (for instance, see the red solid curve around *x* ≈ 0.7 mm in Fig. 3 (a)). As the wave passes, the PVS area increases in a spatial region spanning almost the wavelength of the SD wave, reaching a maximum value of 3.96 × 10^−3^ mm^2^, which is an increase by 69.2% from the baseline value of PVS area.

In Fig. 3 (b), we have plotted the dimensional volumetric flow rate *Q* as a function of *x* using the same convention as Fig. 3 (a). As the wave passes through, the volumetric flow rate increases and CSF flows in the +*x* direction. The increase is in a spatial region spanning almost the wavelength of the SD wave, reaching a maximum value of 7.34×10^−3^ *μ*l/min. The constriction of area induced by the trailing edge of the SD wave leads to a small amount of back flow in the *x* direction of magnitude |*Q*| = 3.23 × 10^−3^ *μ*l/min. Figure 3 (c) shows the variation in the dimensional pressure *P* as the SD wave travels through the PVS segment. In this case, the peak pressure in the PVS as the wave passes through is 6.75 × 10^−3^ mmHg.

We next define a few variables which are useful to quantify the dependence of CSF flow on arterial radius, PVS width, angle of incidence of the SD wave, and the PVS length. We define ⟨*Q*⟩ as the average volumetric flow rate, which is calculated by averaging *Q* along the spatial extent of the PVS, *L*_PVS_, and for a period of *t*_0_ = (*L*_PVS_*/λ* + 1) *T*, which is the time it takes for the SD wave to enter and completely exit a domain of length *L*_PVS_. To characterize the variation of pressure, we define Δ*P*_max_ as the maximum value (during time *t*_0_) of the instantaneous peak pressure difference Δ*P*. We calculate Δ*P* by subtracting the peak and trough pressure values at each instance of time (see Fig. 3 (c)).

### 3.1 The effect of arterial radius and PVS width on CSF flow

We quantify the effect of lumen radius and PVS width on volume flow rate and pressure using the variable Γ which is the ratio of the area of the 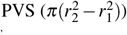 to the area of the artery 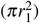. This reduces to 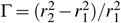 or in terms of PVS width and arterial radius,

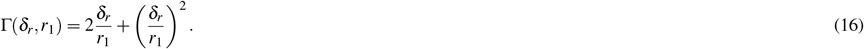

The brain vasculature consists of vessels of variable radii, PVS width, and length. In all the results that follow, we have varied Γ in the range of 0.5≤ Γ≤ 5. Any value of Γ < 0.5 leads to nonphysical results because the PVS width approaches zero/negative values at the constriction induced at the trailing edge of the SD wave. Previous studies have estimated an upper bound for the area ratio as Γ = 2^5,9,59^; however we have numerically probed larger PVS area ratios to gain further insights into the problem.

Figure 4 (a) shows the variation of average nondimensional volume flow rate ⟨*Q*^∗^⟩ as a function of Γ. The simulations are shown by the red data points where ⟨ *Q*^∗^⟩ decreases monotonically with Γ. The solid line is a almost an inverse quadratic power-law curve fit of the form ⟨*Q*^∗^⟩ = *α*_*Q*_/Γ^*η*^_*Q*_, where *α*_*Q*_ = 2.76 and *η*_*Q*_ = 1.9. The inset in the figure shows the same data plotted on a log-log scale. Overall, the fit is excellent. The average dimensional volume flow rate can be obtained by multiplying *Q*_scale_ to the above curve-fit,

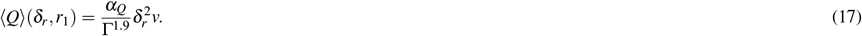

**Figure 4.**
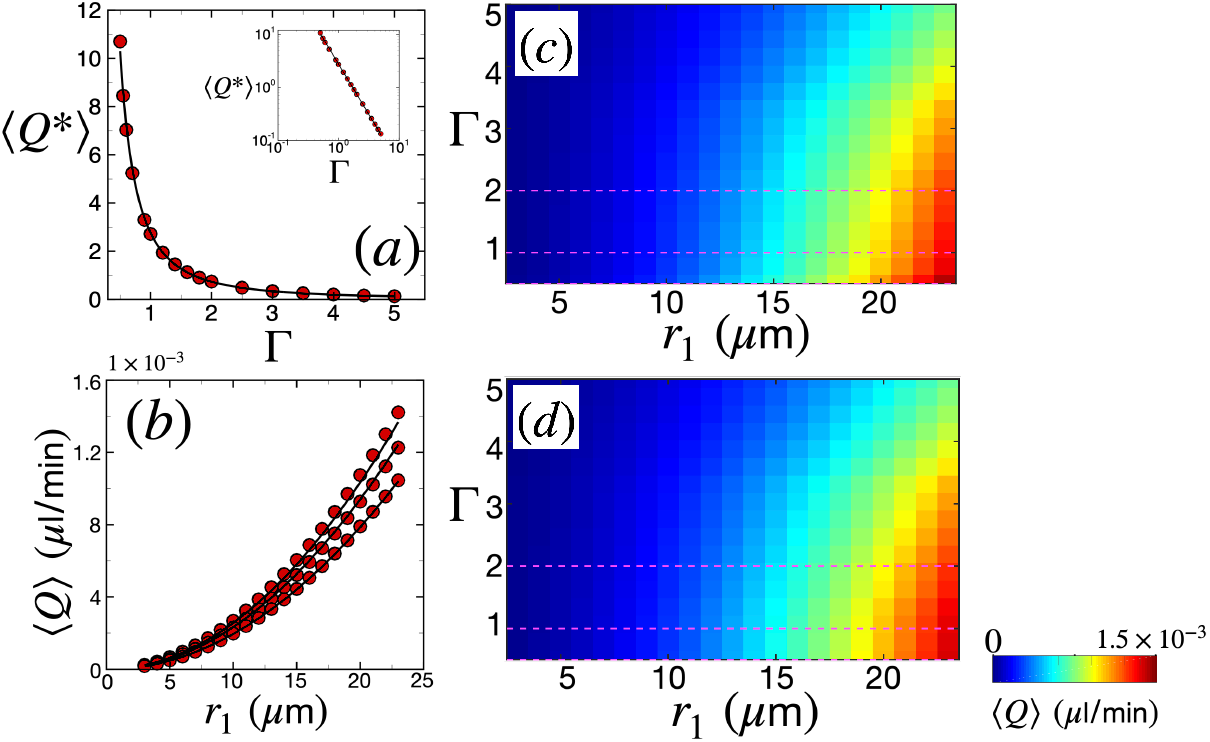
The variation of the spatiotemporally-averaged volume flow rate of CSF ⟨*Q*⟩ as a function of arterial lumen radius *r*_1_ and the width of the PVS quantified using the PVS area ratio Γ (Eq. (16)). Other parameters are *L*_PVS_ = 6 mm and *θ* = 0. (a) Average nondimensional volume flow rate ⟨*Q*^∗^⟩ as a function of Γ. The red circular data points are from simulations and the black solid line is a curve fit through the data of the form ⟨*Q*^∗^⟩ = *α*_*Q*_/Γ^*η*^_*Q*_, where *α*_*Q*_ = 2.76 and *η*_*Q*_ = 1.9. The inset is the same plot in log-log scale. (b) ⟨*Q*⟩ as a function of arterial lumen radius *r*_1_. From top to bottom in the figure, the data points are for Γ = 0.5, 1, and 2 (shown by the dashed lines in (b)) respectively. The black solid lines are curve-fits through the data using the relation Eq. (17). (c) Color-map of average dimensional volumetric flow rate ⟨*Q*⟩ from simulations. The dashed lines are Γ values corresponding to (b). (d) Color-map of ⟨*Q*⟩ from the relation Eq. (17). The dashed lines are Γ values corresponding to the black solid curves in (b).

Figure 4 (b) shows the variation of average dimensional volumetric flow rate *Q* as a function of *r*_1_. The data points are from the simulations while the black solid line is from the expression in Eq. (17), for different values of Γ (Γ = 0.5, 1, and 2, from top to bottom in Fig. 4 (b)). We have varied the arterial lumen radius in the range of 3 *μ*m ≤ *r*_1_ ≤ 23 *μ*m, which covers a range of vasculature radii from pre-capillaries (thinnest) to pial arteries (thickest)^60–62^. It is important to point out however, that PVSs surrounding the capillaries (*r*_1_ < 5.5 *μ*m^61^) are not expected to deform following SD, since capillaries do not possess smooth muscle cells necessary for contraction in the presence of excess concentration of potassium ions.

Figure 4 (c) shows a colormap of ⟨*Q*⟩ in the two-dimensional space spanned by Γ and *r*_1_, which is useful for quantifying how the intensity of ⟨*Q*⟩ varies with Γ and *r*_1_. For instance, we find the maximum value of ⟨*Q*⟩ at the right bottom corner of the colormap, which is for the thickest vessel and smallest PVS area ratio. The colormap indicates that for any constant value of *r*_1_, the volume flow rate will increase with decreasing values of Γ. This is because smaller PVS areas under sudden expansion lead to a large peak pressure difference that pulls in CSF. On the other hand, for any constant value of Γ, ⟨*Q*⟩ increases with *r*_1_. This is because the considerable constriction (Fig. 2 (b)) of an artery with a larger lumen will displace a larger volume of CSF. Figure 4 (d) shows the colormap of the analytical expression given by Eq. (17), showing excellent resemblance to the simulations.

We next characterize the variation of maximum peak pressure difference in the PVS during SD, Δ*P*_max_, as a function of Γ and *r*_1_, which is shown in Fig. 5. Figure 5 (a) shows the variation of nondimensional maximum peak pressure difference 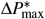 as a function of the PVS area ratio. The data points are from simulations while the solid line is a curve fit. The nature of the data suggests that the peak pressure difference drops monotonically with Γ. Moreover, the drop is steep for Γ < 1 and is more gradual when Γ > 1. This is also evident in the log-log plot in the inset of Fig. 5 (a), where the slope of the data when Γ < 1 is steeper than the data when Γ > 1. We found that the data fits well to a curve which has two power laws describing the two regimes of the form 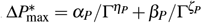, where *α*_*P*_ = 1.19, *β*_*P*_ = 0.4, *η*_*P*_ = 1/2, and *ζ*_*P*_ = 6. The dimensional peak pressure difference can be obtained by multiplying the above relationship by *P*_scale_,

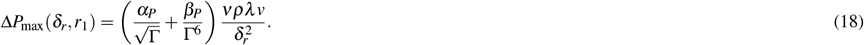

**Figure 5.**
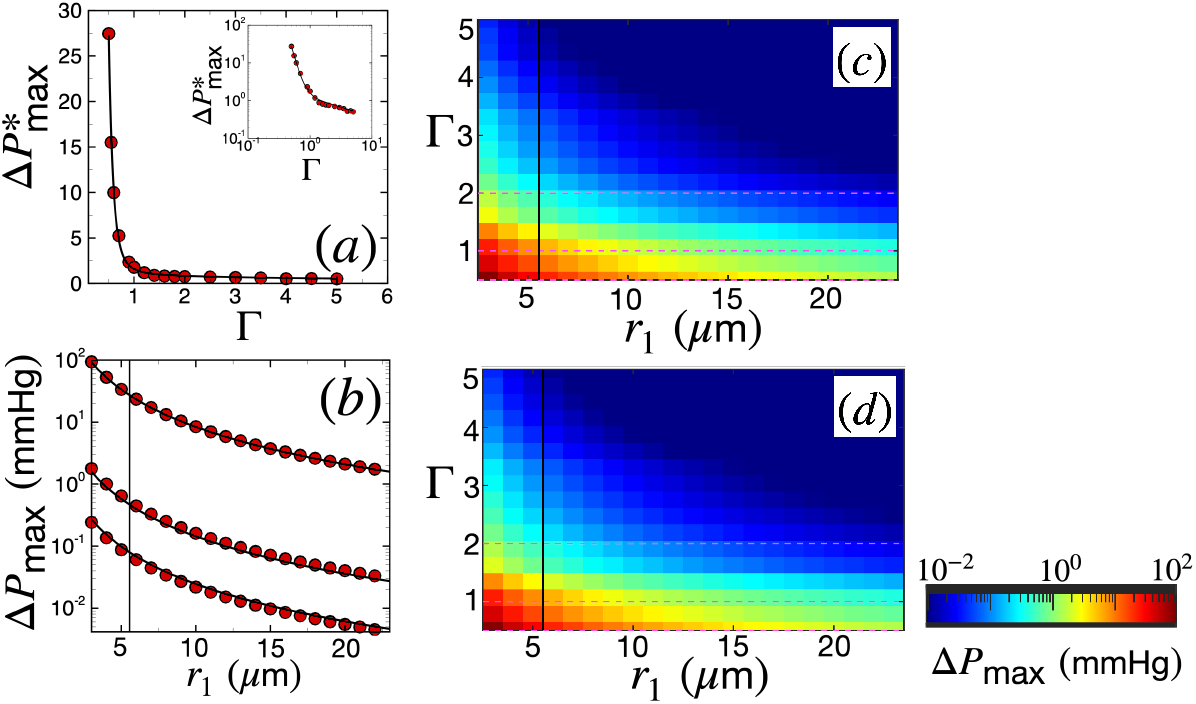
The variation of the maximum peak pressure difference in the PVS during SD as a function of arterial radius *r*_1_ and the width of the PVS quantified using the PVS area ratio Γ (Eq. (16)). Other parameters are *L*_PVS_ = 6 mm and *θ* = 0. (a) Maximum nondimensional peak pressure difference 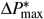 as a function of Γ. The red circular data points are from simulations and the black solid line is a curve fit through the data of the form 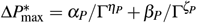, where *α*_*P*_ = 1.19, *β*_*P*_ = 0.4, *η*_*P*_ = 1/2, and *ζ*_*P*_ = 6. The inset is the same plot on a log-log scale. (b) Δ*P*_max_ as a function of arterial radius *r*_1_. From top to bottom in the figure, the data points are for Γ = 0.5, 1, and 2 (shown by the dashed lines in (c)), respectively. The black solid lines are curve fits through the data using the relation Eq. (18). (c) Colormap of dimensional pressure Δ*P*_max_ from simulations. The horizontal dashed lines are Γ values corresponding to the data in (c). (d) Colormap of dimensional peak pressure difference Δ*P*_max_ from the relation in Eq. (18). The horizontal dashed lines are Γ values corresponding to the black solid curves in (b). The data in (c) and (d) is represented by colors that scale logarithmically. The solid vertical line in (b), (c), and (d) is the lower limit for penetrating arteriole radius, *r*_1_ = 5.5 *μ*m, reported in rodents^61^; only results above this limit are expected to be physiologically relevant.

Figure 5 (b) shows the variation of the dimensional maximum peak pressure difference Δ*P*_max_ as a function of the arterial radius *r*_1_ for Γ = 0.5, 1, and 2, from top to bottom in the figure, respectively. The data points from simulations are shown in red and the solid lines are curve fits of the form shown in Eq. (18).

Figure 5 (c) shows a colormap of Δ*P*_max_ in the space of Γ and *r*_1_. The Δ*P*_max_ data is represented by colors that scale logarithmically, with red and blue denoting large and small magnitudes, respectively. The figure shows that maximum peak pressure differences are created when both Γ and *r*_1_ are small (left bottom corner in the figure). As explained before, smaller PVS areas create large peak pressure differences upon sudden expansion. On the other hand, the pressure scales inversely as the square of the vessel radius 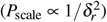, which is why vessels with smaller radius lead to larger pressure differences. Pressure differences reach as high as 94 mmHg in the range of Γ and *r*_1_ that we have explored. However, it is important to note that the lower limit of radius of penetrating arterioles reported in rodents is 5.5 *μ*m^61^. Any vessel with radius lower than this limit can be classified as a pre-capillary, which lacks smooth muscle cells and therefore is not expected to be affected by SD-induced radius alteration. This lower radius limit for vessels affected by SD is indicated by a solid vertical line in Fig. 5 (b), (c), and (d). For *r*_1_ = 5.5 *μ*m, we find a maximum peak pressure difference of about 24 mmHg for Γ = 0.5. Figure 5 (d) shows a colormap of the analytical form shown in Eq. (18) which exhibits excellent agreement with Fig. 5 (c).

### 3.2 The effect of variable angle of incidence of SD waves on CSF flow

An SD wave can approach a pial or a penetrating artery at different angles. We define *θ* as the angle a propagating SD wave is incident on an artery, such that 0≤ *θ* ≤ *π/*2 (see Fig. 1). For an SD wave with an angle of incidence *θ*, the effective wavelength of the wave is *λ*_*θ*_ = *λ/* cos *θ* and the effective velocity of the wave is *v*_*θ*_ = *v/* cos *θ*, whereas the PVS width and arterial radius stay constant. We can predict the volumetric flow rate and pressure variations with *θ* from the expressions derived in the previous section. From Eq. (17) and Eq. (18) the average volumetric flow rate varies with wave velocity and wavelength as⟨*Q*_*θ*_⟩ ∝ *v*_*θ*_ and Δ*P*_max_ ∝ *v*_*θ*_ *λ*_*θ*_, since *r*_1_ and *δ*_*r*_ do not change when the SD wave is incident at an angle. The average volumetric flow rate generated by an SD wave incident at an angle *θ* to the PVS thus reduces to ⟨*Q*_*θ*_⟩ = ⟨*Q*_*θ*=0_⟩ / cos *θ*, where *Q*_*θ*=0_ is the volumetric flow rate when *θ* = 0. Similarly, the maximum peak pressure difference generated due to an SD wave incident at an angle *θ* to the PVS thus reduces to Δ*P*_max,*θ*_ = Δ*P*_max,*θ*=0_/ cos^2^ *θ*, where Δ*P*_max,*θ*=0_ is the maximum peak pressure difference when *θ* = 0.

### 3.3 The effect of PVS length on CSF flow

We next quantify CSF volume flow rate as a function of the length of the PVS, *L*_PVS_. The results discussed so far are for a PVS of length *L*_PVS_ = 2.5*λ* = 6 mm. The brain vasculature consists of vessel segments of variable lengths^61,62^. Here we have explored vessel segments with lengths varying in the range of 0.5 ≤ *L*_PVS_ ≤ 6 mm. Images of mouse cerebral arteries obtained from light sheet microscopy indicate that this range is reasonable (see Fig. 1(I) in Ref.^60^). Additionally, the length of the middle cerebral artery in rats is around 9 mm^70^. Even though precise measurements of the wavelength of SD based on the potassium ion concentration is difficult to obtain experimentally, different numerical models of SD yield a wavelength based on potassium ion concentration of around *λ* = *O*(1) mm^23^. Additionally, the wavelength of SD based on calcium imaging in experiments is *λ* = 𝒪(1) mm (Fig. 3 in Ref.^8^). This implies that SD waves induce fluid flow in both vessels which are shorter and in vessels which are longer than the wavelength of an SD wave. Although the wavelength of SD waves can vary between numerical models and experiments, here we were interested in understanding the variation of CSF dynamics with the ratio of *L*_PVS_*/λ* . In order to understand the effect of domain length on CSF flow rate induced by SD, we explore the variation of ⟨*Q*⟩ with the ratio of domain length to wavelength of the SD wave, *L*_PVS_/*λ* in Fig. 6. Figure 6 (a) shows the variation of average nondimensional volumetric flow rate ⟨*Q*^∗^⟩ as a function of the domain length to wave length ratio *L*_PVS_/*λ* . The data points are from simulations with varying Γ values. We find that ⟨*Q*⟩^∗^ increases monotonically for *L*_PVS_ < *λ*, reaches a maximum value around *L*_PVS_ = *λ* and then decreases monotonically for *L*_PVS_ > *λ* . This suggests that an optimum PVS length for maximizing the volumetric flow rate is *L*_PVS_≈ *λ* .

**Figure 6.**
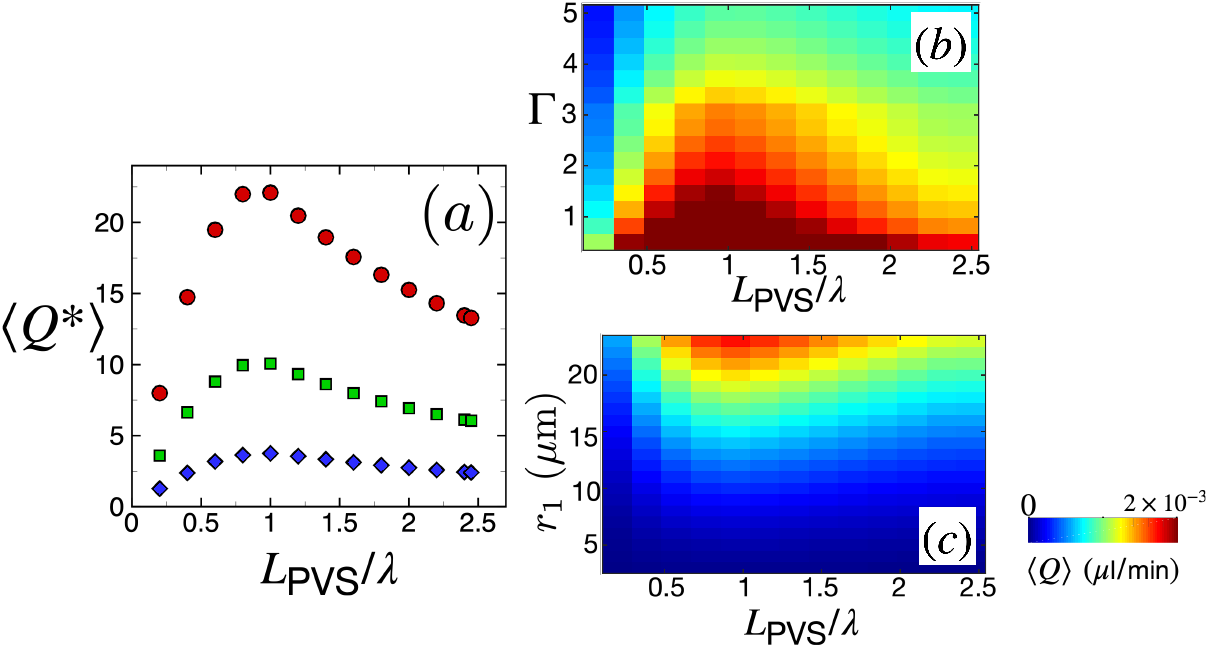
The variation of the CSF volume flow rate with the length of perivascular spaces. (a) Nondimensional average volumetric flow rate *Q*^∗^ as a function of domain length *L*_PVS_. The data points are from simulations with Γ = 0.5 (red, circles), Γ = 0.7 (green, squares), and Γ = 1.1 (blue, diamonds). (b) Colormap of dimensional average volume flow rate *Q* in the space of Γ and domain length *L*_PVS_ for an arterial lumen radius of *r*_1_ = 23 *μ*m. (c) Colormap of dimensional average volume flow rate ⟨*Q*⟩ in the space of arterial lumen radius *r*_1_ and domain length *L*_PVS_. The data corresponds to Γ = 1.5.

Figure 6 (b) shows the colormap of the dimensional average volumetric flow rate ⟨*Q*⟩ in the space of Γ and domain length for a fixed arterial radius of *r*_1_ = 23 *μ*m. The colormap indicates that for any fixed value of domain length, ⟨*Q*⟩ increases as Γ decreases. Additionally, for all values of Γ, the average volume flow rate reaches its maximum when *L*_PVS_ = *λ* .

Figure 6 (c) shows the colormap of the dimensional average volumetric flow rate ⟨*Q*⟩ in the space of *r*_1_ and domain length for a fixed PVS area ratio of Γ = 1.5. The colormap indicates that ⟨*Q*⟩ decreases with arterial lumen radius *r*_1_ for any fixed value of domain length. Moreover, the plot also indicates that volume flow rate is maximized around *L*_PVS_ = *λ* for all values of *r*_1_.

### 3.4 Colliding spreading depolarization waves

SD events typically consist of multiple waves propagating in multiple directions. Indeed, collisions of multiple SD waves among themselves and with the boundaries have been observed in experiments, that lead to complex spatiotemporal dynamics^47,49^. SD waves are known to annihilate each other upon collision^23,49^. Here, we are interested in exploring the variation of CSF volume flow rate and pressure in a PVS when two SD waves collide.

Figure 7 shows the variations in PVS area, CSF volume flow rate, and pressure when two SD waves collide head-on. We simulate these two SD waves propagating in opposite directions, either parallel or anti-parallel to the vessel axis. The first wave is the regular SD wave studied so far which propagates in the +*x* direction and is initiated at *x* = 0. The second wave is an SD wave that propagates in the − *x* direction and is initiated at *x* = *L*_PVS_. The two waves collide at *x* = *L*_PVS_/2. For this study we use an arterial diameter of *r*_1_ = 23 *μ*m, a PVS width of *δ*_*r*_ = 12.63 *μ*m, and an angle of incidence of SD wave equal to *θ* = 0, to drive meaningful comparisons with Fig. 3. It is important to note that similar to our previous results, we have again smoothed out the peaks at the leading edges of the colliding waves due to spatial resolution issues discussed in Sec. 3 and the supplementary information.

**Figure 7.**
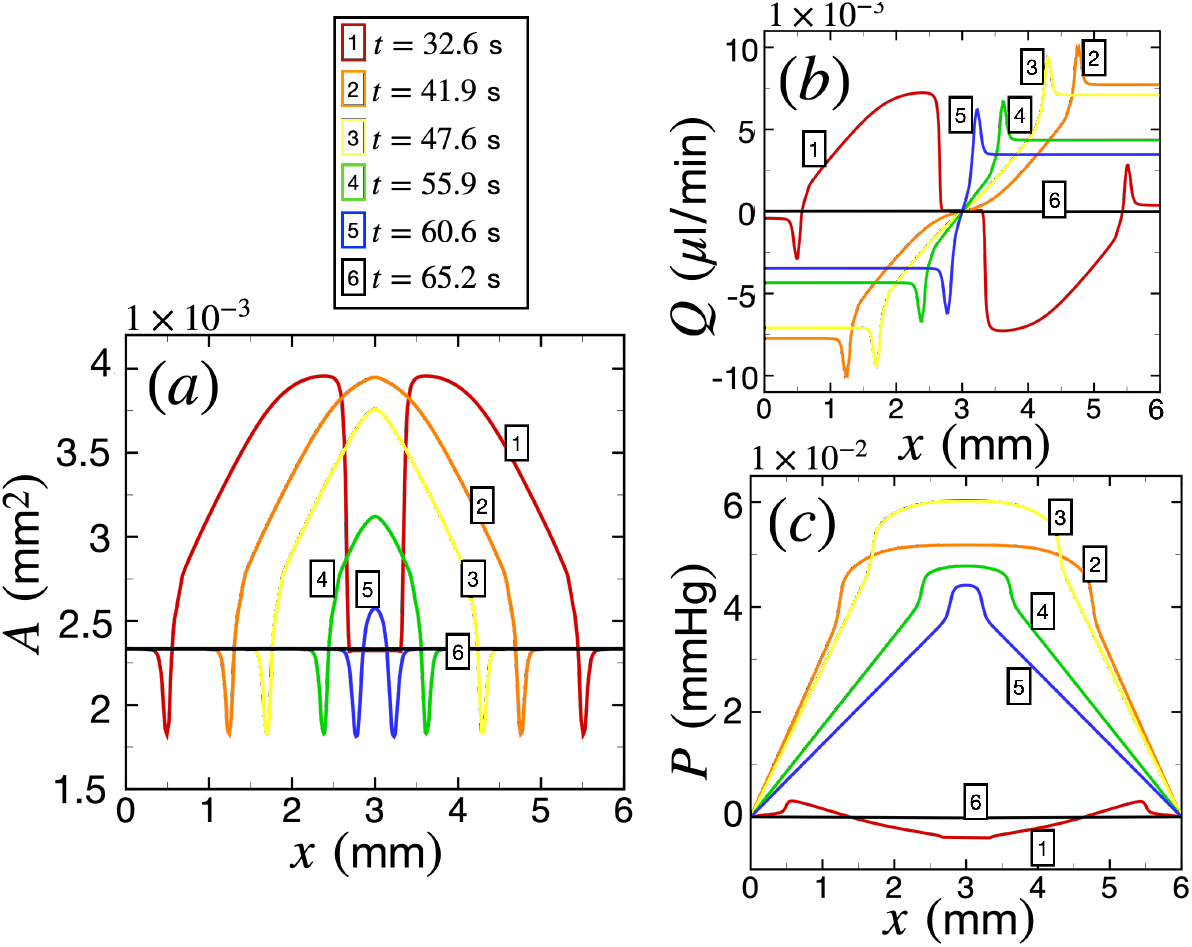
The variation of the area of perivascular space (PVS), volumetric flow rate of cerebrospinal fluid and pressure during a collision of spreading depolarization waves. The arterial lumen radius is *r*_1_ = 23*μ*m, PVS width is *δ*_*r*_ = 12.63*μ*m, and the angle of incidence of the SD waves on the PVS is *θ* = 0. (a) The variation of the area of the PVS, *A* as a function of *x* at different instances of time. The curves at different instances of time are numbered and colored for ease of visualization as *t* = 32.6 s (1, red), *t* = 41.9 s (2, orange), *t* = 47.6 s (3, yellow), *t* = 55.9 s (4, green), *t* = 60.6 s (5, blue), and *t* = 65.2 s (6, black). (b) The variation of volumetric flow rate of cerebrospinal fluid *Q* as a function of *x* using the same plotting convention as (a). (c) The variation of pressure as a function of *x* using the same convention as (a) and (b).

Figure 7 (a) shows the variation of PVS area with space at different instances of time as the two SD waves collide. The different instances of time are numbered in increasing order with time. The two waves start with similar amplitudes as the single-wave case, 3.96 × 10^−3^ mm^2^ (see curves numbered 1, colored red at *t* = 32.6 s). The waves collide at *t* = *L*_PVS_/(2*v*) = 36.2 s. Right after the collision, the waves merge to form a large spatial region of excess potassium ion concentration which acts on the PVS area shown at *t* ≈ 41.9 s (numbered 2, orange). Immediately after this event, the waves annihilate each other as shown by the area decreasing to the baseline value as time progresses (see curves numbered and colored 3, yellow; 4, green; 5, blue; and 6, black).

Figure 7 (b) shows the variation of CSF volume flow rate with space at different instances of time as the two SD waves collide. Before the collision, the absolute value of the maximum volumetric flow rate for both the waves is 7.34 × 10^−3^ *μ*l/min (see the curve numbered 1, colored red), which is similar to Fig. 3 (b). However, right after the collision when the waves merge, *Q* becomes larger attaining a maximum value of 1×10^−2^ *μ*l/min at time *t* = 41.9 s (see the curve numbered 2, colored orange). *Q* then decreases gradually with time as the SD waves annihilate each other and the PVS area decreases to its base value. It is interesting to point out that for this case, a collision of two SD waves lead to an instantaneous *Q* of magnitude 36% larger right after collision than the maximum value of *Q* driven by a single SD wave.

Figure 7 (c) shows the variation of pressure in a single PVS segment due to the collision of two SD waves. Before the collision the peak pressure difference driven by the SD waves is about 7.5 × 10^−3^ mmHg (see the curve numbered 1, colored red). Right after the collision when the waves merge, the peak pressure rises considerably to a maximum value of 0.06 mmHg (see curve numbered 3, yellow). This is a substantial increase in pressure difference, almost 8-fold larger than a single SD wave. In our study, we consider CSF flow in a PVS surrounding a single vessel with the source of CSF flow driven solely by PVS. However, in reality, CSF is already in motion in the brain due to various other mechanisms such as arterial pulsations^9^. In such a scenario, this instantaneous and localized increase in pressure may propel fluid from the PVS into the surrounding brain parenchyma, potentially worsening brain edema.

## 4 Conclusions

We have quantified the coupling of spreading depolarization waves with CSF flow in a PVS lining a single artery for a range of different parameters including length of the PVS, arterial radius, PVS thickness, and the angle of incidence of the SD wave on the PVS. Additionally, we have explored the variation of CSF flow rate and pressure during a collision of two SD waves. Our modeling approach consists of coupling a physiologically realistic spreading depolarization wave in 1D with a 1D Navier-Stokes equation describing the variation in pressure and volume flow rate of CSF due to changes in the cross-sectional area of the PVS. The constriction and dilation of the PVS area is based on an empirical relation tied to excess potassium concentration in the extracellular space (Eq. (15))^35,36^.

We find that an SD wave can lead to substantial CSF flow and pressure gradients in the PVSs of pial and penetrating arteries. Our results show the volume flow rate increases with increasing arterial radius and decreasing PVS width (Fig. 4). We derive an analytical expression of average volume flow rate that agrees well with our simulations in Eq. (17). We also find that the peak pressure difference increases as PVS width and/or arterial radius decrease (Fig. 5). Again, we derive an analytical expression for the maximum peak pressure difference through a PVS with arbitrary width and lumen radius in Eq. (18).

We also quantify the dependence of CSF flow on the length of the PVS (Fig. 6). Our results indicate that CSF flow is maximized around a PVS length which equals the wavelength of the SD wave (*L*_PVS_ = *λ*) for all values of PVS thickness and inner radius. Additionally, we find that the average volumetric flow rate and maximum peak pressure difference depend on the angle of incidence *θ* of the SD wave on the PVS, such that ⟨*Q*_*θ*_⟩ = ⟨*Q*_*θ*=0_⟩ / cos *θ* and Δ*P*_max,*θ*_ = Δ*P*_max,*θ*=0_/ cos^2^ *θ*, where *Q*_*θ*=0_ and Δ*P*_*θ*=0_ are volumetric flow rate and peak pressure differences when *θ* = 0.

Lastly, we explore the alteration in PVS area, CSF volume flow rate, and pressure following a collision of two SD waves (Fig. 7). We find that SD waves annihilate each other upon collision which agrees well with prior studies^23,47,49^. Right after the collision, our simulations indicate that the CSF volume flow rate increases and that SD waves can generate peak pressure differences in the PVSs which are 8-fold larger than the typical pressure differences generated by a solitary SD wave. This excess instantaneous peak pressure difference may drive substantial CSF flow from the PVS, into the brain parenchyma.

There are several limitations in this work that are important to highlight. Firstly, we have used an idealized boundary conditions for the pressure which is *P* = 0 at each end of the PVS. In the future, the effect of different boundary conditions should be more carefully studied. Secondly, although we have explored the variation in CSF flow for a range of PVS thicknesses and vessel radii, we have not considered bifurcations. Bifurcating PVSs may induce different CSF flow response locally at the bifurcation due to an SD wave. A possible future direction is to couple the SD wave with a network model of CSF flow^5,59^.

Our model indicates that SD can induce substantial CSF flow. To compare our results with CSF flow measured in physiological conditions in the murine brain, we refer to the quantitative measurements reported by Mestre et al.^9^. For a typical PVS segment of radius *r*_1_ = 23 *μ*m, length *L*_PVS_ = 6 mm, incident SD angle of *θ* = 0, and area ratio of Γ = 1.4, we obtain an average volume flow rate of ⟨*Q*⟩ = 1.2 ×10^−3^ *μ*l/min which is comparable to the CSF volume flow rate of 2.6× 10^−3^ *μ*l/min measured in Ref.^9^. Additionally, our model indicates that there is an optimum PVS length (equal to the wavelength of the SD wave), which maximizes the CSF volume flow rate following SD. For instance, for the the same parameters outlined above, ⟨*Q*⟩≈ 1.8 × 10^−3^ *μ*l/min when *L*_PVS_ = *λ* . This implies that SD can drive significant CSF flow in the PVS, which may substantially exacerbate brain edema following stroke^8^, cardiac arrest^28^, and traumatic brain injury^29,30^. The analytical expressions that we have derived will be insightful for predicting CSF volume flow rate and pressure changes in a given PVS following SD. For instance, using our results, we can predict regions of the brain which are more susceptible to edema following stroke due to the local orientation of the vasculature. This may, in turn, prove useful in reducing and/or preventing the progression of brain edema.

It is important to point out, however, that a complete picture of edema likely necessitates accounting for the swelling of neurons due to osmotic pressure gradients created by SD^8^. One possible future direction is to investigate how excess CSF flow in the PVSs, driven in via vasoconstriction following SD, subsequently enters the brain parenchyma due to the large osmotic pressure gradients created following SD^33^. This can be achieved by coupling our 1D model to a 2D/3D porous media model of the ECS^10^ with a spatially and temporally-variable extracellular volume fraction *ε*. The value of *ε*, which also appears in Eq. (1-5), could be obtained by modeling the osmotic water influx into the cells following SD. Such a model could then be used to explore severity and prevention strategies for post-SD edema.

Our modeling approach is very general and can be easily adapted to study other spatiotemporal waves in the brain such as slow waves during sleep and functional hyperemia^15,16,63^. Recently, stimulation of cranial nerves (such as the vagus nerve) has been shown to enhance CSF flow^41,64^. Additionally, electrical stimulation creates local ion sinks which potentially may drive CSF flow via a mechanism similar to SD. Our model can help design such neural stimulation protocols. For instance, using our model one can predict CSF flow through PVSs lining vasculature adjacent to the nerve that is stimulated. Lastly, our results form a foundation for investigating localized augmentation of CSF flow in the brain by controlling SD, which may prove therapeutically beneficial. SD can be initiated in a controlled way through targeted neuromodulation of brain vasculature using externally applied electrical fields^65^. One could leverage this approach to initiate SD in a localized part of the brain with large amyloid-*β* buildup (linked with Alzheimer’s disease), potentially aiding its removal through the SD-associated CSF flow enhancement.

## Author contributions statement

J.T. and S.M. conceived of the project. S.M. developed and performed the modeling and numerical simulations of SD. M.M. performed the modeling and simulations of the 1D Navier-Stokes equations. S.M. and M.M. performed the modeling and data analysis for coupled SD and CSF flow. All authors contributed to the interpretation of data. S.M. wrote the initial draft of the manuscript. All authors revised the text and approved of the final version.

## Data availability

All data used in this paper is available on request. The corresponding author S.M. should be contacted for requesting the data from this study.

