## Supplementary Information for "Quantifying the relationship between spreading depolarization and perivascular cerebrospinal fluid flow"

### 1 Parameters used in spreading depolarization simulation

The initial resting values of ions and gating variables used in the SD simulation are shown in Supplementary Table S(1). The ionic currents and gating variables in the SD model governed by the Goldman-Hodgkin-Katz equation is shown in Supplementary Table S(2). Supplementary Table S(3) shows the conductances used for Goldman-Hodgkin-Katz and Hodgkin-Huxley leak currents, respectively.

| Parameter | Values |
|---|---|
| $[\mathrm{K}]_e$ | 3.86 mM |
| $[\mathrm{K}]_i$ | 133.45 mM |
| $[\mathrm{Na}]_e$ | 141.6 mM |
| $[\mathrm{Na}]_i$ | 9.82 mM |
| $m_{\mathrm{NaT}}$ | 9.82 |
| $h_{\mathrm{NaT}}$ | 0.995 |
| $m_{\mathrm{NaP}}$ | $1.3\times10^{-2}$ |
| $h_{\mathrm{NaP}}$ | 0.1 |
| $m_{\mathrm{KDR}}$ | $1.3\times10^{-3}$ |
| $h_{\mathrm{KDR}}$ | 0.1 |
| $m_{\mathrm{KA}}$ | 0.12 |
| $h_{\mathrm{KA}}$ | 0.12 |

**Supplementary Table S(1).** Initial resting values of ions and gating variables used in the SD simulation taken from Ref.[1].

| Currents (mA/cm$^2$) | Gates, $m^P h^q$ | Rate Constants |
|---|---|---|
| $I_{\text{Na},T}$ | $m^3h$ | $\alpha_m = \frac{0.32(E_m+51.9)}{1-e^{-(0.25E_m+12.975)}}$<br>$\beta_m = \frac{0.28(E_m+24.89)}{e^{-(0.2E_m+4.978)}-1}$<br>$\alpha_h = 0.128e^{-(0.056E_m+2.94)}$<br>$\beta_h = \frac{4}{1+e^{-(0.2E_m+6)}}$ |
| $I_{\text{Na},P}$ | $m^2h$ | $\alpha_m = \frac{1}{6(1+e^{-[0.143E_m+5.67]})}$<br>$\beta_m = \frac{e^{-(0.143E_m+5.67)}}{6(1+e^{-(0.143E_m+5.67)})}$<br>$\alpha_h = 5.12\times10^{-8}e^{-(0.056E_m+2.94)}$<br>$\beta_h = \frac{1.6\times10^{-6}}{1+e^{-(0.2E_m+8)}}$ |
| $I_{\text{K},DR}$ | $m^2$ | $\alpha_m = \frac{0.016(E_m+34.9)}{1-e^{-(0.2E_m+6.98)}}$<br>$\beta_m = 0.25e^{-(0.025E_m+1.25)}$ |
| $I_{\text{K},A}$ | $m^2h$ | $\alpha_m = \frac{0.02(E_m+56.9)}{1-e^{-(0.1E_m+5.69)}}$<br>$\beta_m = \frac{0.0175(E_m+29.9)}{e^{(0.1E_m+2.99)}-1}$<br>$\alpha_h = 0.016e^{-(0.056E_m+4.61)}$<br>$\beta_h = \frac{0.5}{1+e^{-(0.2E_m+11.98)}}$ |

**Supplementary Table S(2).** Gating variables and rate constants for active currents governed by the Goldman-Hodgkin-Katz equation used in the SD simulation following Ref.[1].

| Currents (mA/cm$^2$) | $g_{\text{ion},GHK}$ | $g_{\text{ion},HH}$ |
|---|---|---|
| $I_{\text{Na},T}$ | $100\times10^{-5}$ | |
| $I_{\text{Na},P}$ | $2\times10^{-5}$ | |
| $I_{\text{K},DR}$ | $100\times10^{-5}$ | |
| $I_{\text{K},A}$ | $10\times10^{-5}$ | |
| $I_{\text{Na,leak}}$ | | $2\times10^{-5}$ |
| $I_{\text{K,leak}}$ | | $7\times10^{-5}$ |
| $I_{\text{leak}}$ | | $20\times10^{-5}$ |

**Supplementary Table S(3).** Conductances of active GHK currents and passive HH leak currents used in the SD simulation following Ref.[1].

### 2 Derivation of 1D Navier-Stokes equation

To simulate the effect of SD on the CSF flow, the change in arterial radius is calculated based on the potassium ion concentration from the SD simulations and then fed into the reduced Navier-Stokes equations. To this end, a 1D formulation of the Navier-Stokes equations for an incompressible fluid flow between two concentric cylinders is derived. Assuming an axisymmetric flow with axial and radial velocity components, the governing continuity and momentum equations in cylindrical coordinates are[2]:

$$\frac{1}{r}\frac{\partial(ru_r)}{\partial r} + \frac{\partial u_x}{\partial x} = 0, \tag{1}$$

$$\frac{\partial u_r}{\partial t} + u_r\frac{\partial u_r}{\partial r} + u_x\frac{\partial u_r}{\partial x} + \frac{1}{\rho}\frac{\partial P}{\partial r} = \nu\left(\frac{1}{r}\frac{\partial}{\partial r}(r\frac{\partial u_r}{\partial r}) + \frac{\partial^2 u_r}{\partial x^2} - \frac{u_r}{r^2}\right), \tag{2}$$

$$\frac{\partial u_x}{\partial t} + u_r\frac{\partial u_x}{\partial r} + u_x\frac{\partial u_x}{\partial x} + \frac{1}{\rho}\frac{\partial P}{\partial x} = \nu\left(\frac{1}{r}\frac{\partial}{\partial r}(r\frac{\partial u_x}{\partial r}) + \frac{\partial^2 u_x}{\partial x^2}\right), \tag{3}$$

where $u_r$ and $u_x$ are fluid velocity components in the radial and axial directions, $P$ is pressure, $\rho$ is the fluid density, $\nu$ is the kinematic viscosity of the fluid, $t$ is time, and $x$ and $r$ are the axial and radial coordinates. To derive the reduced 1D

Navier-Stokes equations, Eqs. (1)-(3) are non-dimensionalized and averaged in the radial direction. To non-dimensionalize these equations we use the following transformations:

$$r^* = r/\delta_r, \tag{4}$$

$$u_r^* = \frac{u_r \lambda}{v \delta_r}, \tag{5}$$

$$u_x^* = u_x/v, \tag{6}$$

$$x^* = x/\lambda, \tag{7}$$

$$t^* = t/\tau, \tag{8}$$

$$P^* = P\frac{\delta_r^2}{\mu \lambda v}, \tag{9}$$

where $\delta_r = r_2 - r_1$ is the initial thickness of the annulus, $\lambda$, $\tau$, and $v$ are respectively the wavelength, temporal period, and wave speed of the motion of the arterial wall as a function of potassium ion concentration, and $\mu$ is the dynamic viscosity of CSF. The Reynolds number obtained from such nondimensionalization is:

$$\text{Re} = \frac{v{\delta_r}^2}{\nu \lambda}. \tag{10}$$

Using the nondimensionalized parameters and defining an aspect ratio $\omega = \delta_r/\lambda$, Eq. (1), Eq. (2), and Eq. (3) can be written as:

$$\frac{1}{r^*}\frac{\partial(r^* u_r^*)}{\partial r^*} + \frac{\partial u_x^*}{\partial x^*} = 0, \tag{11}$$

$$\text{Re}\ \omega^2\left(\frac{\partial u_r^*}{\partial t^*} + u_r^*\frac{\partial u_r^*}{\partial x^*} + u_x^*\frac{\partial u_r^*}{\partial x^*}\right) + \frac{\partial P^*}{\partial r^*} = \omega^2\left(\frac{1}{r^*}\frac{\partial}{\partial r^*}(r^*\frac{\partial u_r^*}{\partial r^*}) + \omega^2\frac{\partial^2 u_r^*}{\partial x^{*2}} - \frac{u_r^*}{r^{*2}}\right), \tag{12}$$

$$\text{Re}\left(\frac{\partial u_x^*}{\partial t^*} + u_r^*\frac{\partial u_x^*}{\partial r^*} + u_x^*\frac{\partial u_x^*}{\partial x^*}\right) + \frac{\partial P^*}{\partial x^*} = \left(\frac{1}{r^*}\frac{\partial}{\partial r^*}(r^*\frac{\partial u_x^*}{\partial r^*}) + \omega^2\frac{\partial^2 u_x^*}{\partial x^{*2}}\right). \tag{13}$$

In our model $\omega^2 \ll 1$, which means the lubrication approximation will be valid. As a result, Eq. (12) and Eq. (13) are reduced to:

$$\frac{\partial P^*}{\partial r^*} = 0, \tag{14}$$

$$\text{Re}\left(\frac{\partial u_x^*}{\partial t^*} + u_r^*\frac{\partial u_x^*}{\partial r^*} + u_x^*\frac{\partial u_x^*}{\partial x^*}\right) + \frac{\partial P^*}{\partial x^*} = \frac{1}{r^*}\frac{\partial}{\partial r^*}\left(r^*\frac{\partial u_x^*}{\partial r^*}\right). \tag{15}$$

Equation (14) indicates that pressure variation along the radial direction is negligible. Furthermore, Eq. (11) and Eq. (15) can be manipulated into:

$$\frac{\partial(r^* u_r^*)}{\partial r^*} + \frac{\partial(r^* u_x^*)}{\partial x^*} = 0 \tag{16}$$

$$\text{Re}\left(\frac{\partial(r^* u_x^*)}{\partial t^*} + \frac{\partial(r^* u_r^* u_x^*)}{\partial r^*} + \frac{\partial(r^* u_x^{*2})}{\partial x^*}\right) + \frac{\partial(r^* P^*)}{\partial x^*} = \frac{\partial}{\partial r^*}\left(r^*\frac{\partial u_x^*}{\partial r^*}\right). \tag{17}$$

Assuming a Poiseuille flow velocity profile, the axial fluid velocity is[2]:

$$u_x^* = -\frac{1}{4}\frac{\partial P^*}{\partial x^*}\left[r_2^{*2} - r^{*2} + \left(\frac{r_2^{*2} - r_1^{*2}}{\log(\frac{r_1^*}{r_2^*})}\right)\log\left(\frac{r_2^*}{r^*}\right)\right]. \tag{18}$$

By integrating Eq. (16) and Eq. (17) in the radial direction, then using the boundary condition $[u_r]_{r=r_1} = \partial r_1/\partial t$ at the arterial wall, we obtain:

$$\frac{\partial A^*}{\partial t^*} + \frac{\partial Q^*}{\partial x^*} = 0, \tag{19}$$

$$\mathrm{Re}\left(\frac{\partial Q^*}{\partial t^*}+\frac{\partial(Q^*\Psi_u u^*)}{\partial x^*}\right)+A^*\frac{\partial P^*}{\partial x^*}=-4c^*Q^*, \tag{20}$$

in which the following definitions are used:

$$Q^*=A^*U^*, \tag{21}$$

$$A^*=\pi(r_2^{*2}-r_1^{*2}), \tag{22}$$

$$U^*=\frac{1}{A^*}\int_{r_1^*}^{r_2^*}2\pi u_x^* r^* dr^*, \tag{23}$$

$$\Psi_u=\frac{1}{u^{*2}A^*}\int_{r_1^*}^{r_2^*}2\pi u_x^{*2} r^* dr^*. \tag{24}$$

$\Psi_u$ is the fraction of the square of the horizontal velocity to the square of the average velocity and $Q^*$ is the averaged volumetric flow rate. The parameter $c^*$ is given by:

$$c^*=\frac{2}{r_2^{*2}+r_1^{*2}+\frac{r_2^{*2}-r_1^{*2}}{log(r_1^*/r_2^*)}}. \tag{25}$$

Since $\mathrm{Re}=O(10^{-3})$, we neglect the nonlinear term inside the parentheses of Eq. (20). However, we retain the first term since the problem is time-dependent. The 1D reduced equations for fluid flow that we then solve are,

$$\frac{\partial A^*}{\partial t^*}+\frac{\partial Q^*}{\partial x^*} = 0, \tag{26}$$

$$\mathrm{Re}\frac{\partial Q^*}{\partial t^*}+A^*\frac{\partial P^*}{\partial x^*} = -4c^*Q^*. \tag{27}$$

We implement a predictor-corrector method to numerically solve the equations. The predictor step predicts a value of $Q^*$ using the momentum equation given by Eq. (27). We then take the divergence of Eq. (27) and use the mass equation given by Eq. (26) to obtain a Poisson equation in pressure. After solving the Poisson equation, we correct the value of $Q^*$ and march forward in time. The predictor step is:

$$\frac{\delta Q^*}{\delta t^*}=\frac{1}{\mathrm{Re}}\left(-4c^*Q^*-A^*\frac{\delta P^*}{\delta x^*}\right). \tag{28}$$

Taking the divergence of Eq. (27) and writing the equation in finite differences yields:

$$\frac{\delta^2 P^*}{\delta x^{*2}}=\frac{1}{A^*}\left(-4Q^*\frac{\delta c^*}{\delta x^*}+4c^*\frac{\delta A^*}{\delta t^*}-\frac{\delta A^*}{\delta x^*}\frac{\delta P^*}{\delta x^*}+\mathrm{Re}\frac{\delta^2 A^*}{\delta t^{*2}}\right). \tag{29}$$

We solve the Poisson equation using the method of direct inversion of the tridiagonal matrix. After obtaining a corrected pressure field, we use that field to correct the $Q^*$ field in the corrector step by again solving Eq. (28). We use a 4th order accurate explicit Runge-Kutta method in time and 2nd order accurate finite differences in space. The numerical method is implemented in Fortran.

### 3 Relationship between arterial radius and potassium ion concentration

Supplementary Figure S(1) shows a plot of the normalized arterial lumen radius as a function of the extracellular potassium ion concentration $[\mathrm{K}]_e$ following Eq. (15) in the main text, adopted from Refs.[3,4].

### 4 Accuracy of the numerical approach

For testing the accuracy of our numerical approach we have conducted convergence tests of the SD and 1D Navier-Stokes solvers separately. Supplementary Figure S(2) shows the spatial convergence of the SD and 1D Navier-Stokes solvers. Supplementary Figure S(2) (a) shows the spatial convergence of the relative error $\epsilon_v$ as a function of the nondimensional spatial resolution $\Delta x^*=\Delta x/\lambda$ for the SD simulations conducted using $\Delta t=0.005$ ms. Since there are no exact analytical solutions, the relative error is computed using the wave speed at the finest resolution $\Delta x=0.002$ mm that we could simulate for this time step.

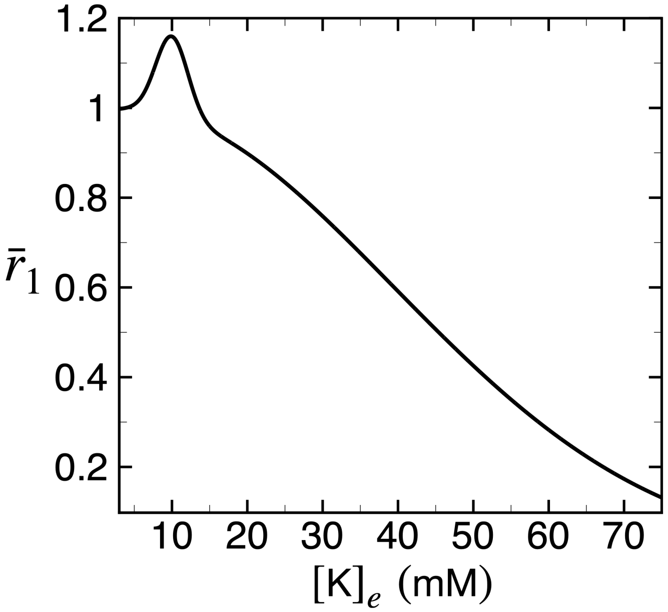


**Supplementary Figure S(1).** The variation of the normalized arterial lumen radius as a function of the extracellular potassium ion concentration $[K]_e$ using the empirical relationship given by Eq. (15) in the main text, adopted from Refs.[3,4].

We then plot $\epsilon_v = (v - v_{\text{finest}})/v_{\text{finest}}$ as a function of grid resolution. Since we employ a 2nd order accurate in space finite difference approach, we expect the data to follow a line of slope 2 when plotted on a log-log scale. Indeed, we find that with finer grid resolutions the data approaches a line of slope 2 on the log-log plot. However, it is challenging to conclusively address the accuracy of a numerical approach in the absence of an exact solution. Since discretization errors in finite difference schemes that are 2nd order accurate in space are expected to follow $\epsilon_v \propto \Delta x^2$, we can also find the order of accuracy by comparing $\epsilon_v$ at a particular grid resolution $P = \Delta x$ and $q = \Delta x/2$. The order of accuracy can then be estimated by the quantity $\gamma = \log(\epsilon_{v,P}/\epsilon_{v,q})/\log 2$. For instance, when $P = 0.01$ mm and $q = 0.005$ mm, we have $\gamma = 2.2$ in our 2nd order accurate code. We find that the value of $\gamma$ varies within 15% of the expected order of accuracy for the range of grid spacings we have explored. We used an explicit 4th order accurate Runge-Kutta method in time and found the largest possible time step we could implement to be $\Delta t = 0.005$ ms.

Supplementary Figure S(2) (b) shows the spatial convergence of the 1D Navier-Stokes solver. To analyze the spatial convergence we have coupled a sine wave of fixed amplitude and wavelength with the 1D Navier-Stokes equations, for which an approximate analytical solution exists[5]. We solve Eq. (A10) in Ref.[5] numerically using Newton's iterative method to obtain the analytical solution. We then compare our spatiotemporally averaged volumetric flow rate with the analytical solution and plot the relative error $\epsilon_Q = \left(\langle Q\rangle - \langle Q\rangle_{\text{analytical}}\right)/\langle Q\rangle_{\text{analytical}}$ as a function of grid resolution. The black solid line in Supplementary Fig. S(2) (b) has a slope of 2 which is expected for a 2nd order accurate spatial finite difference formulation. We used an explicit 4th order accurate Runge-Kutta method in time and found that we had to satisfy a Courant–Friedrichs–Lewy condition of $\Delta t^*/\Delta x^* < 0.03$.

### 5 Effect of the sharp peak in radius at the leading edge of the SD wave on the results

The raw data of radius variation as a function of $x$, induced by the SD wave, includes peaks at both leading and the trailing edges of the radius distribution (Fig. 2(b) in the main text). Supplementary Figure S(3) (a) shows the variation of the PVS area induced by the SD wave, where the red, green, and blue square data points are at different times in the evolution of the wave. Note that the peaks in radius in the former plot correspond to troughs in PVS area in the latter plot. Figure S(3) (a) shows that the dips in PVS area (i.e., values less than about $2.4 \times 10^{-3}$ mm$^2$) occur abruptly, and the trough at the leading edge is particularly sharp (indicated by the very few data points that make up the right trough). The grid in this case is unable to resolve this sharp feature. The effect of the under-resolved trough is shown in Supplementary Figure S(3) (b), where we plot the variation of $Q$ at different instances of time. The presence of the under-resolved trough leads to unrealistic oscillations in the volumetric flow rate (as well as the pressure, which is not shown). Our grid spacing for solving the fluid flow is typically $\Delta x = 0.01$ mm. A straightforward approach can be to use a larger number of grid points to resolve the feature. However, we found that we are not able to resolve the leading sharp peak even after decreasing the grid spacing to $\Delta x = 0.001$ mm,

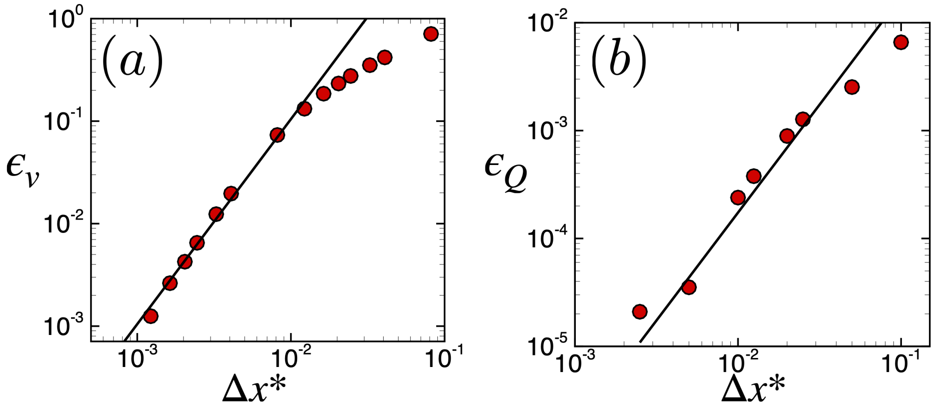


**Supplementary Figure S(2).** Plots of relative error as a function of spatial resolution for the SD and 1D Navier-Stokes simulations respectively. (a) Relative error of SD wave velocity $\epsilon_v = (v - v_{\text{finest}})/v_{\text{finest}}$ as a function of $\Delta x^* = \Delta x/\lambda$, where $\epsilon_v$ is computed with respect to the wave velocity at the finest resolution. (b) Plot of relative error of volumetric flow rate $\epsilon_Q = \left(\langle Q\rangle - \langle Q\rangle_{\text{analytical}}\right)/\langle Q\rangle_{\text{analytical}}$ as a function of $\Delta x^*$, where $\epsilon_Q$ is computed with respect to the analytical solution of $Q$ obtained from Ref.[5] In both (a) and (b), the black solid lines have a slope of 2 which is expected from the 2nd order accurate spatial finite difference formulation.

which came at a substantial computational cost. In addition to requiring extremely small time steps to resolve the fine grid spacing, we also reach a computational bottleneck when trying to solve the Poisson equation in pressure (Eq. (29)) using direct inversion. In terms of the results, the case with non-smoothed variations in PVS area yields an average volumetric flow rate of $\langle Q\rangle = 0.00125$ $\mu$l/min, while a smoothed case that removes the leading trough in PVS area leads to $\langle Q\rangle = 0.00117$ $\mu$l/min while also minimizing unrealistic oscillations, and hence the effect of smoothing out this feature is small.

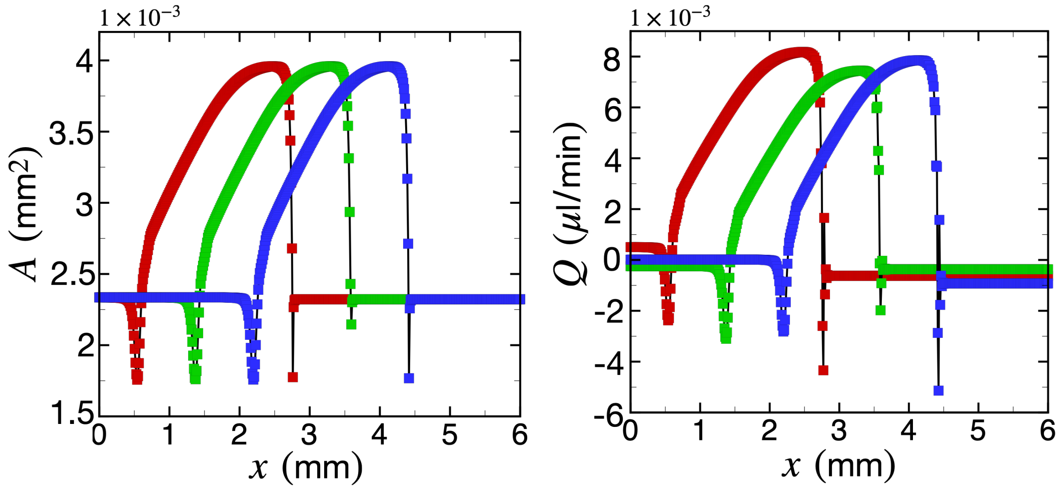


**Supplementary Figure S(3).** Non-smoothed data of variation of (a) PVS area and (b) CSF volumetric flow rate in a PVS subjected to an SD wave under the same parameters studied in Fig. 3 of the main manuscript.
